# A Genetically Encoded Actuator Selectively Boosts L-type Calcium Channels in Diverse Physiological Settings

**DOI:** 10.1101/2023.09.22.558856

**Authors:** Pedro J. del Rivero Morfin, Diego Scala Chavez, Srinidhi Jayaraman, Lin Yang, Audrey L. Kochiss, Henry M. Colecraft, X. Shawn Liu, Steven O. Marx, Anjali M. Rajadhyaksha, Manu Ben-Johny

**Author notes:** To whom correspondence must be addressed: Manu Ben-Johny, PhD.

## Abstract

L-type Ca^2+^ channels (Ca_V_1.2/1.3) convey influx of calcium ions (Ca^2+^) that orchestrate a bevy of biological responses including muscle contraction and gene transcription. Deficits in Ca_V_1 function play a vital role in cardiac and neurodevelopmental disorders. Yet conventional pharmacological approaches to upregulate Ca_V_1 are limited, as excessive Ca^2+^ influx leads to cytotoxicity. Here, we develop a genetically encoded enhancer of Ca_V_1.2/1.3 channels (GeeC) to manipulate Ca^2+^ entry in distinct physiological settings. Specifically, we functionalized a nanobody that targets the Ca_V_ macromolecular complex by attaching a minimal effector domain from a Ca_V_ enhancer—leucine rich repeat containing protein 10 (Lrrc10). In cardiomyocytes, GeeC evoked a 3-fold increase in L-type current amplitude. In neurons, GeeC augmented excitation-transcription (E-T) coupling. In all, GeeC represents a powerful strategy to boost Ca_V_1.2/1.3 function in distinct physiological settings and, in so doing, lays the groundwork to illuminate new insights on neuronal and cardiac physiology and disease.

## INTRODUCTION

L-type calcium (Ca^2+^) channels (Ca_V_1.2 and Ca_V_1.3) are biologically crucial as they convert electrical signals to Ca^2+^ influx that orchestrates an impressive array of physiological processes^1-3^. In the heart, Ca_V_1.2 participates in shaping the action potential (AP), and in initiating excitation-contraction coupling^4^. During adrenergic stimulation, increased Ca_V_1.2 activity leads to improved inotropy, resulting in enhanced heart contraction and ejection fraction^5^. In neurons, both Ca_V_1.2 and Ca_V_1.3 channels impact neuronal excitability through interplay with Ca^2+^-activated K^+^ channels ^6,7^, and support excitation-transcription coupling that contributes to synaptic plasticity^8-11^. Beyond this, Ca_V_1.2 and/or Ca_V_1.3 channels play important roles in various physiological settings including: (1) tuning insulin secretion in pancreatic beta cells^12^, (2) regulating AP firing and catecholamine release from adrenal chromaffin cells^13^, (3) smooth muscle contraction tuned Ca^2+^-entry through Ca_V_1.2^14^, (3) vesicle secretion from ribbon synapses in inner ear hair cells that relies exclusively on presynaptic Ca_V_1.3^15^, and (4) varied emerging functions in non-excitable cells^16^. Alterations in Ca_V_1.2 and Ca_V_1.3 are, therefore, linked to diverse clinical phenotypes (see reviews^17-19^). Heterozygous gain-of-function Ca_V_1.2 variants are linked to Timothy Syndrome, a multisystem disorder characterized by prolonged QT interval, syndactyly, and autism spectrum disorders (ASD)^20^. On the other hand, loss-of-function Ca_V_1.2 variants are associated with Brugada Syndrome, a cardiac arrhythmogenic disorder where the cardiac AP is shortened, resulting in short QT interval and increased likelihood for sudden death^21^. Furthermore, recent studies have identified loss-of-function Ca_V_1.2 variants linked to variable neurological phenotypes including developmental delays, ASD, and affective disorders^21,22^. As such, strategies to upregulate Ca_V_1.2/1.3 function may be therapeutically beneficial^23^. Pharmacologically, Ca_V_1 blockers including dihydropyridines and phenylalkylamine are used clinically for various indications^2,24^. However, activation of Ca_V_1 channels is a double-edged sword—non-tissue specific pharmacological Ca_V_1 agonists can lead to broad activation of Ca_V_1 channels resulting in cytotoxicity. In rodents, Ca_V_1 specific agonists Bay K 8644 and FPL 64176 result in severe motor abnormalities^25^, seizures^26^, and self-harming behavior^27^. Therefore, alternate strategies for targeted or tissue specific upregulation of Ca_V_1 function are highly sought after^28^.

Beyond this, both Ca_V_1.2 (*CACNA1C*) and Ca_V_1.3 (*CACNA1D*) have been found to be prominent risk genes for various neurodevelopmental and neuropsychiatric disorders (see reviews^29,30^) including ASD^20^, schizophrenia^31,32^, and bipolar disorder^33^, suggesting important yet complex functions in the brain. Even still, the fundamental neurobiological mechanisms that contribute to these varied phenotypes remain to be fully elucidated. The emerging consensus is that either upregulation or reduction in channel function can contribute to human disease, likely depending on the specific cell-type or neural circuit involved^29^. However, our current understanding of the physiological functions of these channels and their pathophysiological consequences are derived largely from either constitutive or cell-type specific knockout of Ca_V_1.2 and/or Ca_V_1.3 in mice^3,34^. By comparison, there are limited strategies available to upregulate Ca_V_1.2/1.3 function in a cell-type dependent manner or with subcellular specificity. This technological limitation hampers an in-depth understanding of biological mechanisms. From an experimental perspective, development of next-generation customizable Ca_V_1 actuators are highly advantageous to illuminate their variegated physiological functions.

Here, we sought to develop a molecular actuator of Ca_V_1.2/Ca_V_1.3 activity that upregulates channel activity in distinct physiological settings. To this end, beyond the pore-forming subunit that is targeted by conventional pharmacology^35^, Ca_V_ channels associate with an impressive array of regulatory proteins^36^, several of which interact with channel cytosolic domains and finely tunes channel dynamics^2,37^. We reasoned that leveraging an intracellular regulatory protein may provide an alternative strategy to engineer a Ca_V_ enhancer. In this regard, we utilized a recently identified cardiac-specific Ca_V_1.2 modulator leucine rich repeat containing protein 10 (Lrrc10) that upregulates Ca_V_1.2 current in heterologous systems and ventricular cardiomyocytes^38^. Lrrc10 has been implicated in cardiomyocyte maturation^39,40^ and regeneration^41-43^ as well as in the cardiac response to increased afterload^44^. We identified a minimal effector domain within Lrrc10 that upregulates Ca_V_1.2, albeit with a low affinity such that it is unable to modulate channel function by itself. Instead, fusion of this domain to a nanobody that binds the Ca_V_ channel complex with a high affinity yielded a Ca_V_1.2/1.3 specific actuator termed—genetically encoded enhancer of Ca^2+^ channels (GeeC)—that markedly increases the channel open probability (*P*_O_). We further deployed ^45^GeeC into cardiomyocytes and neurons to modulate endogenous Ca_V_1.2/Ca_V_1.3 current and tune downstream physiological functions.

## RESULTS

### Leveraging Lrrc10 modulation to engineer a Ca_V_1.2 actuator

To quantify the effect of Lrrc10 on Ca^2+^ currents, we undertook whole cell recordings of Ca_V_1.2, co-transfected with auxiliary subunits α_2_δ and β_2b_ in HEK293 cells and measured peak current density. When compared to cells expressing only Ca_V_1.2 canonical subunits (Figure 1A), co-expression of Lrrc10 yielded a nearly 5-fold increase in peak current density (Figure 1B), consistent with previous reports^38^. Given this striking increase in current density, we sought to identify key segments within Lrrc10 responsible for binding and modulating Ca_V_ channels. Similar to other leucine rich repeat (Lrr) containing proteins, Lrrc10 is composed of seven stereotypic Lrr domains with signature sequence LxxLxLxxNxL or LxxLxLxxNxxL flanked by short N-terminal (NT) and C-terminal (CT) regions^46^. Traditionally, the Lrr domains are thought to enable binding to disparate targets, allowing this diverse family of proteins to serve a wide-range of cellular functions^47^. We used a flow cytometry-based FRET 2-hybrid assay to probe interaction of Lrrc10 segments with holo Ca_V_1.2 in live cells (Figure 1C). Specifically, we bisected Lrrc10 into (1) an N-terminal (NT) domain and (2) a C-terminal (CT) segment that included the Lrr domains and attached to cerulean fluorescent protein, the FRET donor, to each peptide. Venus fluorescent protein, the FRET acceptor, was attached to the carboxy terminus of Ca_V_1.2. Indeed, full length Lrrc10 displayed robust FRET binding with holo-Ca_V_1.2 (Figure 1C). Furthermore, the Lrrc10 CT segment also revealed strong binding to Ca_V_1.2 with a relative association constant (*K*_a,EFF_) that is nearly identical to full-length Lrrc10 (Figure 1C). By comparison, the isolated Lrrc10 NT domain showed weak to no-binding (Figure1C). These results suggest that the Lrrc10 CT containing the Lrr domains is critical for robust Ca_V_1.2 interaction.

**Figure 1.**
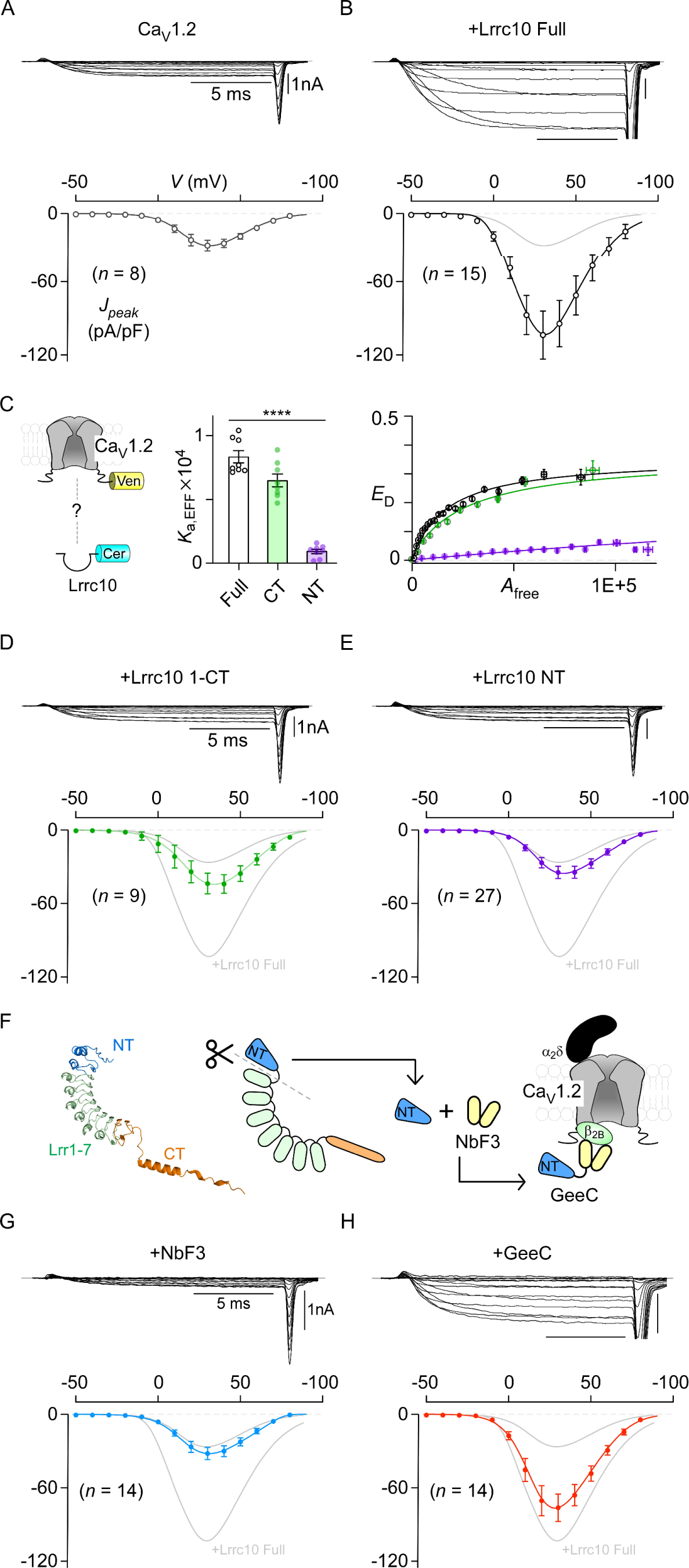
Leveraging Lrrc10 modulation of Ca_V_1.2 channels to develop a genetically encoded enhancer. **(A)** Whole cell recordings show robust baseline Ca_V_1.2 currents in HEK293 cells. *Top*, exemplar traces show Ca_V_1.2 current evoked by 15 ms depolarizations to a family of test pulse potentials. *Bottom*, aggregate data for current density-voltage (*I*-*V*) relationship. n, number of cells. Error bars show mean and s.e.m. **(B)** Co-expression of full-length LRRC10 markedly increases Ca_V_1.2 current density. Format as in panel A. **(C)** *Left.* Cartoon illustrates a FRET pair with Cerulean-tagged Lrrc10 or subsegments serving as FRET donor while Venus-tagged holoCa_V_1.2 serving as FRET acceptor. *Right,* Flow-cytometry based live cell FRET 2-hybrid assay shows that Ca_V_1.2 interacts robustly with full length Lrrc10. Further dissecting Lrrc10 revealed that the CT domain containing the Lrr domains suffice for binding to Ca_V_1.2, while NT domain (gray) exhibits poor binding. **** p < 0.0001. **(D)** Although Lrrc10 CT suffices for strong interaction with Ca_V_1.2, this domain is insufficient for functional channel modulation. Format as in format A. **(E)** Lrrc10 NT also fails to alter Ca_V_1.2 currents. **(F)** *Right*. Cartoon shows structure-guided design of GeeC by fusing the Lrrc10 NT with nb.F3 that targets Ca_V_1.2 **(G)** Co-expression of nb.F3 minimally perturbs peak Ca_V_1.2 current density. Format as in A. **(H)** GeeC markedly enhances peak L-type current density in HEK 293 cells. Format as in A.

Thus affirmed, we probed whether Lrrc10 CT also sufficed for functional modulation of Ca_V_1.2. Surprisingly, we found that co-expression of Lrrc10 CT only minimally changed Ca_V_1.2 peak current density, despite its ability to interact strongly with full length Ca_V_1.2 (Figure 1D). Furthermore, co-expression of the Lrrc10 NT domain also revealed little to no change in peak current density, consistent with its weak affinity for Ca_V_1.2 (Figure 1E). Taken together, these results point to two distinct possibilities: (1) the overall tertiary structure of Lrrc10 may be essential for functional Ca_V_1.2 modulation such that any alteration in its structure abrogates its function, or (2) Lrrc10 may be composed of two functionally distinct domains: the CT segment containing the Lrr domains may be responsible for high affinity binding to Ca_V_1.2, while the NT segment may be a low affinity effector domain responsible for driving Ca^2+^ current upregulation.

We reasoned that if Lrrc10 CT serves as a binding domain for Ca_V_1.2, then replacing this region with a peptide or a domain that interacts with Ca_V_1.2 would preserve functional modulation. In this regard, recent studies have developed a nanobody that binds to the Ca_V_β-subunit, nb.F3, with a high affinity (∼10 nM) but is functionally inert^48^. As such we engineered a chimeric protein by fusing Lrrc10 NT region with nb.F3 and probed its effect on tuning Ca_V_1.2 currents (Figure 1F). Indeed, co-expression of nb.F3 with Ca_V_1.2 yielded no change in peak current density, consistent with the nanobody being functionally silent (Figure 1G). However, co-expression of the chimeric Lrrc10-NT fused to nb.F3 resulted in a nearly 3-fold increase in peak current density (Figure 1H). Given the ability to upregulate Ca_V_1.2 currents, we named this chimeric protein “genetically encoded enhancer of Ca^2+^ currents” or GeeC (ˈgēk). Overall, these findings are consistent with the bi-modular architecture of Lrrc10 whereby the NT serves as a low affinity effector domain to increase currents while the CT serves as a binding domain that is responsible for interacting with Ca_V_1.2. Of note, as Lrr repeats often interacts promiscuously with protein targets, this chimeric approach provides a potential platform for targeted upregulation of Ca_V_ channels in distinct physiological settings.

### GeeC selectively upregulates Ca_V_1.2/1.3 channels

Having confirmed the ability of GeeC to enhance Ca_V_1.2 currents, we sought to determine the selectivity of GeeC in tuning various members of the Ca_V_ channel superfamily. All members of the Ca_V_1/Ca_V_2 family are known to interact with the Ca_V_β subunit targeted by GeeC^49^. As such, it is possible that any of these channels may be modulated by GeeC. By comparison, Ca_V_3 channels do not require the Ca_V_β subunit for its function^50^ and is unlikely to be affected by GeeC. Accordingly, we systematically quantified changes in peak current density of other Ca_V_ channels upon co-expression with GeeC. Similar to Ca_V_1.2, we found that GeeC yielded a 5-fold increase in Ca_V_1.3 currents compared to co-expression of nb.F3 (Figure 2A,2F). By contrast, the closely related Ca_V_1.4 channels were entirely unaffected (Figure 2B, 2F). Furthermore, we found that GeeC had minimal to no effect on Ca_V_2 channels (Figure 2C-2F). These findings demonstrate that GeeC is selective in upregulating Ca_V_1.2/1.3 channels.

**Figure 2.**
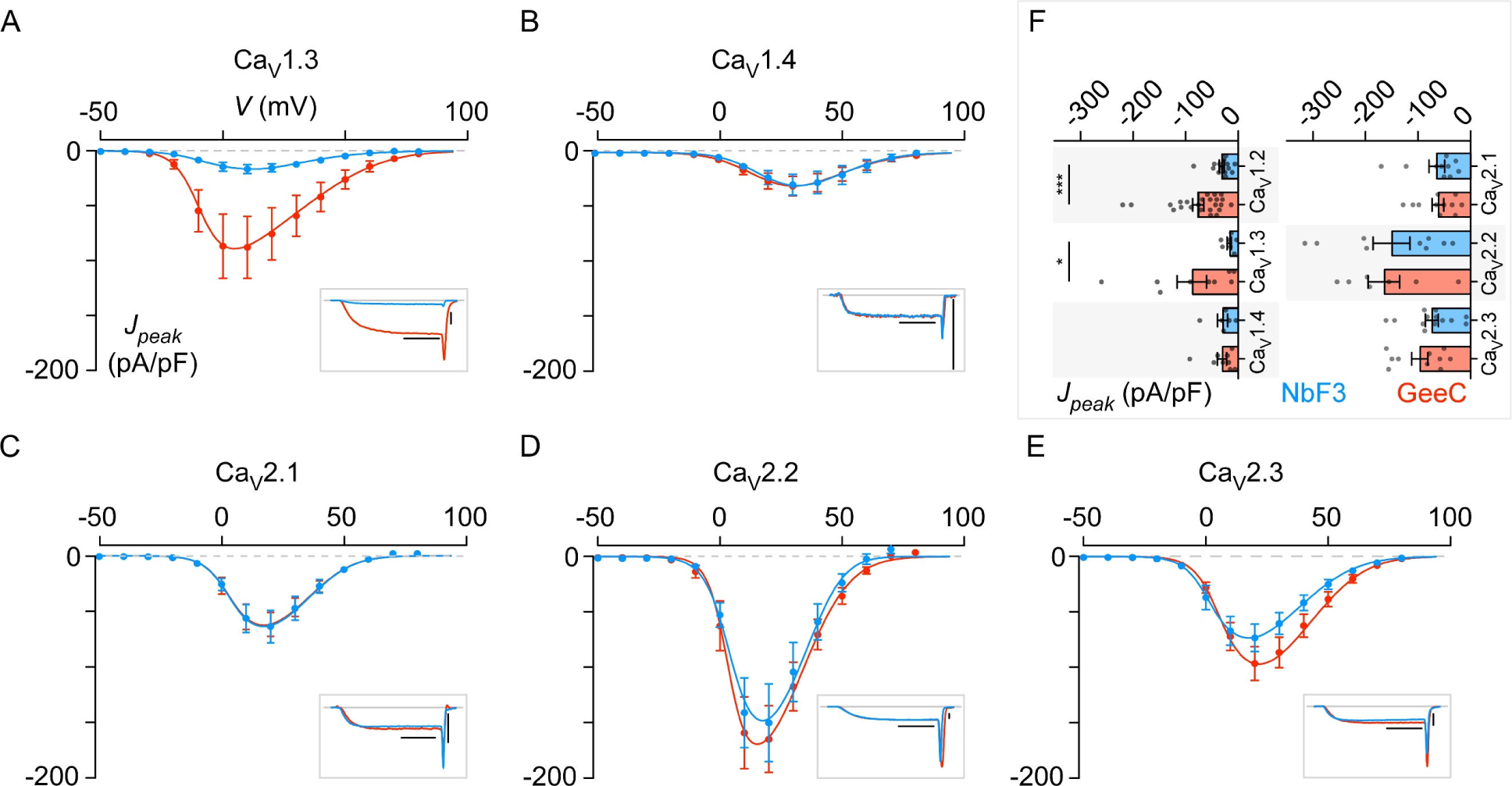
GeeC selectively enhances Ca_V_1.2/1.3 currents. **(A)** Co-expression of GeeC (red) with Ca_V_1.3 subunits increases L-type currents in HEK 293 cells compared to cells co-expressing nb.F3 by itself (blue). Inset shows exemplar Ca_V_1.2 recordings in the presence of nb.F3 or GeeC. Error bars show mean and s.e.m. **(B-E)** GeeC fail to alter peak current density of other closely related Ca_V_ channels. Format as in A. **(F)** Bar graphs compare peak current density for various Ca_V_1/2 channels in the presence of nb.F3 (blue) or GeeC (red). Error bars show mean and s.e.m. * p < 0.05, *** p < 0.001.

### Mechanism of Ca_V_1.2/1.3 upregulation by GeeC

Mechanistically, GeeC may upregulate Ca_V_1.2/1.3 whole cell current by enhancing one of three fundamental elementary channel properties: (1) the open probability (*P*_O_), i.e. the likelihood that channel is open at a given voltage, (2) the unitary current (*i*) that describes the current through a single open channel, or (3) the total number of channels (*N*) at the surface membrane. To distinguish between these possibilities, we undertook low-noise cell attached single-channel recordings of Ca_V_1.2 in the presence of either GeeC or nb.F3 in HEK293 cells (Figure 3A,B). These recordings permit direct measurement of both *P*_O_ and *i*, while preserving the intracellular milieu. Here, we use Ba^2+^ as charge carrier to avoid confounding effects of Ca^2+^-dependent inactivation. Stochastic channel openings were evoked at near steady-state *P*_O_ at each voltage using a slow voltage-ramp. Exemplar recordings in Figure 3A show channel openings as downward deflections to the unitary current level at each voltage (gray slanted curve). Qualitatively, we found that GeeC increased the number and duration of openings in comparison to nb.F3, without perturbing the unitary current, suggesting that GeeC enhances *P*_O_. The steady-state P_O_ – voltage relationship was then estimated by averaging 80 – 150 stochastic records to obtain a mean current that is divided into the open level, and averaged over multiple patches. Thus probed, we found that GeeC increased the maximal *P*_O_ by approximately 3-fold (Figure 3B).

**Figure 3.**
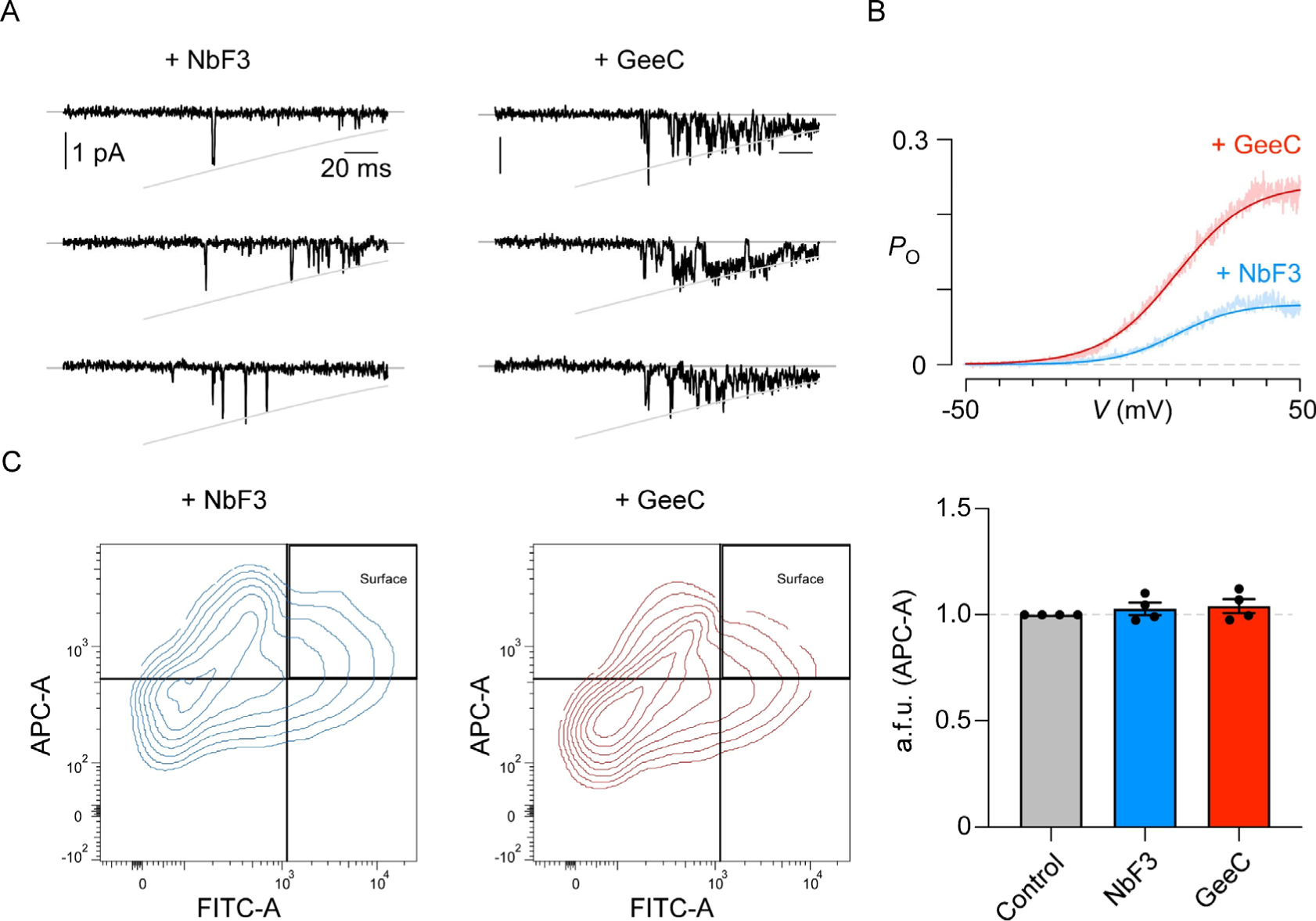
GeeC enhances Ca_V_1.2/1.3 function by upregulating channel openings. **(A)** Exemplar single-channel recordings of Ca_V_1.2 heterologous expressed in HEK293 cells in the presence of either nb.F3 or GeeC. Slanted gray curve represents unitary conductance. Each downward deflection indicate channel openings **(B)** Ensemble average *P*_O_ - voltage relationship for Ca_V_1.2 shows that GeeC increases peak *P*_O_ compared to cells expressing nb.F3 alone. n = 4 (nb.F3) and 5 (GeeC) **(C)** *Left,* exemplar datasets show that nb.F3 and GeeC minimally perturb to Ca_V_1.2 surface membrane trafficking. *Right*, population data comparing surface-membrane expression of Ca_V_1.2 in the presence of either nb.F3 (blue) or GeeC (red). Bars show mean and s.e.m.

To determine whether GeeC alters surface-membrane trafficking of Ca_V_1.2, we used a dual labeling approach that is previously established^51^. Specifically, the pore-forming α-subunit of Ca_V_1.2 was engineered to contain tandem bungarotoxin (BTX) binding sites on a loop exposed to the extracellular surface, and a yellow fluorescent protein (YFP) on its carboxy-terminus. With this assay, the total number of channels in single cells can be measured as the fluorescence intensity of YFP, while the surface-membrane channels can be selectively labeled by incubating cells with BTX conjugated to Alexa-647. GeeC minimally altered surface-membrane trafficking of Ca_V_1.2 channels as reported by normalized Alexa-647 signal (Figure 3C). Taken together, these results suggest that GeeC enhances Ca_V_1.2 function by exclusively boosting channel openings.

### GeeC enhances L-type current in cardiomyocytes

Having established the functionality of GeeC, we sought to determine whether GeeC can upregulate native L-type current in cardiomyocytes. We cultured ventricular cardiomyocytes from adult mice, which primarily express Ca_V_1.2 channels and adenovirally transduced GeeC along with an mCherry fluorescent reporter (Figure 4A). Whole cell current recordings show baseline Ca^2+^ currents in these cells evoked in response to a family of voltage step depolarization (Figure 4B). Adenoviral transduction of green fluorescent protein (GFP) revealed a minor reduction in peak current density (Figure 4C). By contrast, GeeC expression yielded up to 3-fold increase in Ca^2+^ current density (Figure 4D). Importantly, expression of nb.F3 alone failed to upregulate the Ca^2+^ current density, with peak current density matching levels observed with GFP expression (Figure 4E). These findings demonstrate the exquisite ability of GeeC to enhance native Ca_V_1.2 channels in cardiomyocytes.

**Figure 4.**
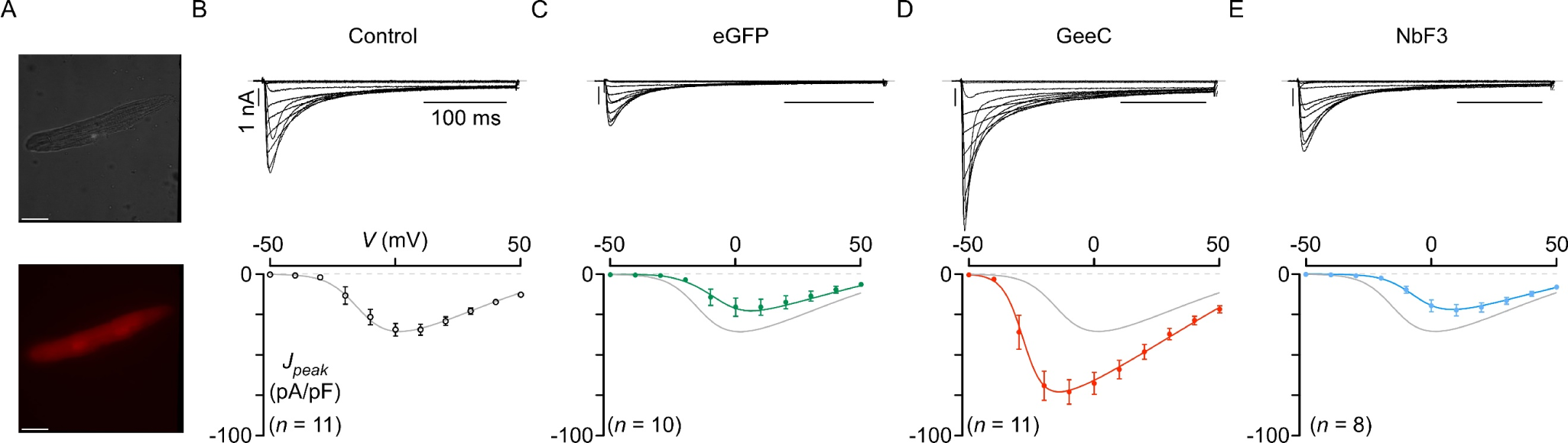
GeeC enhances endogenous Ca_V_1.2 in mouse-derived cardiomyocytes. **(A)** Brightfield (top) and epifluorescence (bottom) images show a cardiomyocyte expressing GeeC, with mCherry bicistronically expressed. **(B)** *Top*, exemplar traces of endogenous L-type currents in 2-day cultured cardiomyocytes evoked in response to 300 ms voltage steps to various potentials. *Bottom*, population data show baseline peak current density. Error bar, mean and s.e.m. *n* = 11 from 3 mice. **(C)** Adenoviral expression of eGFP minimally perturbs L-type current density. *n* = 10 from 3 mice. Format as in panel B. **(D)** GeeC markedly upregulates endogenous L-type channels. *n* = 11 from 3 mice. Format as in panel B. **(E)** Nb.F3 by itself failed to appreciably perturb peak current density. Format as in panel B. n = 8 from 2 mice.

### GeeC boosts excitation-transcription coupling in neurons

In neurons, Ca_V_1.2/1.3 channels play an essential role in coupling excitation to changes in gene expression, a process known as excitation-transcription coupling. This process is critical for driving neurodevelopmental changes and in modulating synaptic plasticity^8,11^. Briefly, Ca^2+^ entry through NMDA and Ca_V_1.2/1.3 channels^9,10^, as well as voltage-dependent conformation changes in the latter^52^, leads to autophosphorylation of Ca^2+^/calmodulin-dependent protein kinase II (CaMKII)^8^. This, in turn, results in phosphorylation and nuclear translocation of immediate early genes such as cyclic AMP-response element binding protein (CREB)^10^. We, therefore, hypothesized that upregulation of Ca_V_1.2/1.3 by GeeC may boost excitation-transcription coupling. To test this possibility, we transduced either GeeC or eGFP into cultured rat hippocampal neurons. As in previous studies, neurons were depolarized with high extracellular potassium (K^+^) for ∼2 mins. The strength of excitation-transcription coupling was then measured by immunostaining of nuclear phosphorylated CREB (pCREB). Without depolarization, minimal pCREB is observed in the nucleus (Figure 5A, 5D). Following depolarization, we observed a significant pCREB staining (∼ 5-fold) in the nucleus (Figure 5A,5D). Adenoviral expression of GFP resulted in minimal changes in depolarization-induced pCREB in the nucleus (Figure 5B, 5D). By comparison, expression of GeeC resulted in a nearly 2-fold increase in nuclear pCREB staining (Figure 5C-5D). These findings reveal the ability of GeeC to augment excitation-transcription coupling in cultured neurons, likely reflecting an enhancement in Ca_V_1.2/1.3 function.

**Figure 5.**
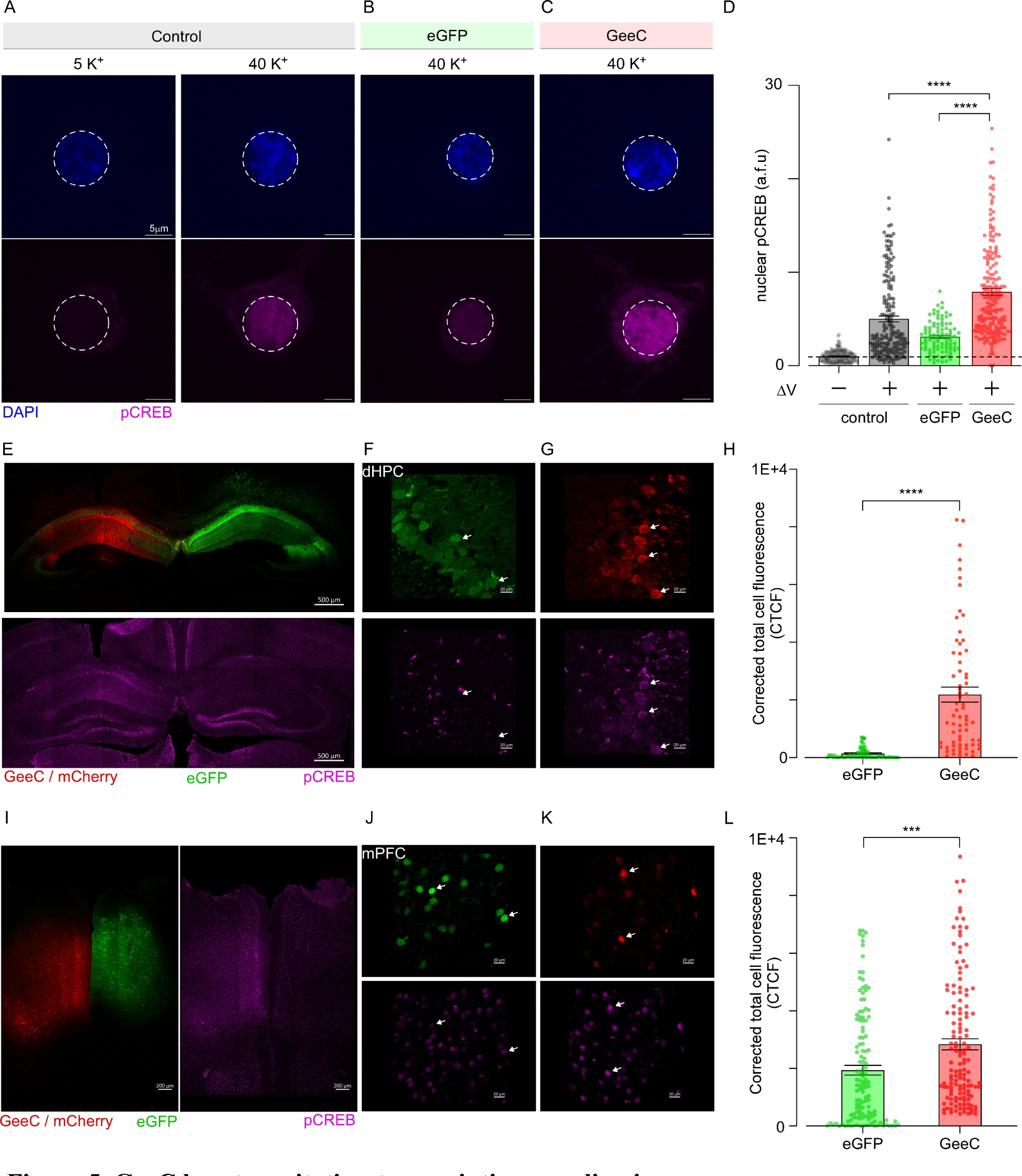
GeeC boosts excitation-transcription coupling in neurons. **(A)** Confocal images increased nuclear accumulation of pCREB follow cell depolarization with extracellular solution containing 40 mM K^+^. Top, DAPI staining; Bottom, pCREB staining (Alexa647). White dashed circles mark the nuclear region of interest for fluorescence quantification. **(B)** Expression of eGFP has minimal effect on nuclear pCREB levels. **(C)** Expression of GeeC qualitatively increases nuclear pCREB staining. **(D)** Population data compares nuclear pCREB signal following neural excitation. Bars show mean ± s.e.m. **** p < 0.0001. **(E)** To probe functionality of GeeC in vivo, we stereotactically injected AAV9 encoding either eGFP or GeeC / mCherry. Exemplar coronal slice of mouse hippocampi at 10x magnification shows robust expression of GeeC (mCherry) or eGFP control AAV9 (top), and pCREB signal (bottom). pCREB staining is qualitatively higher in the right hemisphere with GeeC expression. **(F-G)** Higher magnification (40X) exemplar confocal images of mouse dorsal hippocampal neurons showing mcherry and pCREB or GFP and pCREB. White arrows indicate infected cells. **(H)** Population data shows increased pCREB in GeeC-infected hippocampal neurons. **(I)** Representative coronal slice of mouse prefrontal cortex at 10x magnification shows either GeeC or eGFP transduction (left), and pCREB staining (right). **(J-K)** Higher magnification (40X) exemplar confocal images of mice prefrontal cortex neurons. White arrows indicate infected cells. **(L)** Population data confirms enhanced pCREB staining in GeeC-transduced prefrontal cortex neurons when compared to eGFP-infected controls. Format as in D. *** p < 0.001.

Having confirmed its effect in cultured neurons, we set out to determine whether GeeC can boost excitation-transcription coupling in the mouse brain. Using stereotaxic surgical techniques, we injected unilaterally in the dorsal hippocampus and the medial prefrontal cortex (mPFC) of 12-14 week old mice with either AAV9-GeeC (right hemisphere) or AAV9-eGFP (left hemisphere) as negative control. Following 2.5 weeks of recovery during which the mice were allowed to behave normally, we confirmed robust expression of GeeC (mCherry marker) and GFP in the two hemispheres and probed pCREB expression using immunostaining in brain sections using confocal microscopy. We found that hippocampal neurons from brain hemispheres injected with AAV9-GeeC showed increased pCREB activity compared to hemispheres injected with AAV9-eGFP control (Figure 5E-H). Similar effects were also observed in mPFC neurons, with GeeC expressing neurons demonstrating an increase in pCREB staining compared to GFP control (Figure 5I-L). In all, these findings illustrate the exquisite capability of GeeC to modulate excitation-transcription coupling in neurons.

## DISCUSSION

Precise manipulation of Ca_V_1.2/Ca_V_1.3 in distinct physiological settings is crucial for delineating their biological functions and as potential therapeutics^2^. Conventional pharmacological agonists are limited by potential cytotoxicity owing to broad spectrum activation of Ca_V_1 channels. Here, we engineered GeeC as a genetically encoded approach to boost Ca_V_1.2/1.3 function with enhanced specificity in various physiological settings. To do so, we functionalized a nanobody that binds the Ca_V_β subunit by attaching a low affinity effector domain derived from Lrrc10, a cardiac specific Ca_V_1.2 modulator. In depth analysis revealed that GeeC selectively modulates Ca_V_1.2 and Ca_V_1.3 channels while minimally perturbing closely related Ca_V_ channels, by upregulating channel *P*_O_. In cardiomyocytes, GeeC expression results in increased Ca_V_1.2 current density, confirming its functionality in modulating native Ca^2+^ channels. In cultured hippocampal neurons, we found that GeeC expression markedly increased nuclear translocation of pCREB suggesting increased excitation-transcription coupling, likely owing to increased Ca^2+^ influx through Ca_V_1.2/1.3 channels. Complementing these results, we found that GeeC delivery into mouse hippocampus and medial prefrontal cortex increased pCREB in neurons, suggesting potentiated transcriptional coupling *in vivo*. Thus, GeeC provides a new avenue to selectively upregulate Ca_V_1.2/1.3 function in various physiological settings.

Several pharmacological Ca_V_1 blockers including dihydropyridines and verapamil are routinely used clinically to downregulate Ca_V_1 activity for various indications^2^. Although Ca_V_1.2/1.3 agonists have been postulated as potential treatment for multiple diseases including bladder dysfunction^23^, these approaches pose inherent challenges. Broad activation of these channels across multiple tissues will likely result in excessive Ca^2+^ entry, leading to toxicity and off-target effects. Indeed, previous studies that used Bay K 8644, a dihydropyridine Ca_V_1 agonist, in rodents reported increased coronary vascular resistance, self-biting behavior, and characteristic motor abnormalities including twisting and stretching movements, limb extension, back arching, spasticity, and ataxia^25-27^. By contrast, as GeeC is genetically encoded, emerging viral gene delivery approaches^53^ or other protein targeting approaches can be used to deliver GeeC to specific cell types to precisely modulate Ca_V_1.2/1.3 function without evoking potential off-target effects or toxicity. From a biological perspective, GeeC may provide a convenient strategy to dissect the functional contribution of Ca_V_1.2/1.3 channels in various physiological settings. For instance, in the brain, inducible expression of GeeC in distinct neuron populations or neural circuits may allow us to dissect the complex contribution of Ca_V_1 function to elicit a variety of behavioral responses in animal models. Similarly, GeeC may be beneficial to determine how Ca_V_1 channels contribute to the generation of neural circuits in the developing brain through inducible expression in early developmental stages^54^. In the heart, GeeC could be utilized to selectively target Ca_V_1.2 channels in cardiomyocytes, potentially increasing E-C coupling, inotropy and cardiac output. This may be advantageous to better delineate the discrete roles for augmented Ca^2+^ entry versus phosphorylation in heart remodeling and cardiac disease where prolonged adrenergic stimulation plays a central pathophysiological role^55,56^.

Our findings also provide new insights on the mechanism of Lrrc10 modulation of Ca_V_1 channels. Lrrc10 is an evolutionarily conserved protein that is exclusively expressed in cardiomyocytes^57^. In zebrafish, Lrrc10 is essential for cardiac function, including cardiac development^42^. Specifically, Lrrc10 knockdown leads to reduced cardiomyocyte population, and cardiac dysmorphism. In mice, deletion of Lrrc10 results in reduced cardiac function prior to birth that progresses to dilated cardiomyopathy in adulthood^57,58^. In addition, these mice exhibit reduced contractility in response to pressure overload induced by traverse aortic constriction^44^. Furthermore, human Lrrc10 variants are associated with pediatric dilated cardiomyopathy^38,59^ and cardiac arrhythmias^60^. Recently, Lrrc10 has also been found to be an essential element for heart regeneration in both zebrafish^41,42^ and in mammals^43^; however, how Lrrc10 mediates this effect is not fully understood. Lrrc10 is known to associate with multiple proteins, including the α-subunit of the Ca_V_1.2 channel^38^, α-actinin^58^, and actin^58^. Here, we find that the high affinity interaction with Ca_V_1.2 is mediated by the Lrr-domains, similar to Lrrc10 interaction with other targets. Indeed, Lrr domains are thought to mediate protein-protein interactions and may promiscuously support association with diverse targets^46^. Critically, functional Ca_V_1.2 upregulation also requires the NT of Lrrc10, which serves as a low affinity effector domain. Unlike the Lrr domains which share considerable homology with other Lrrc family of proteins^57^, the NT region is unique to Lrrc10. This suggests that while other members of the Lrrc family of proteins may also interact with the Ca_V_1.2 channel, it is unlikely that these proteins would serve as Ca_V_1.2/1.3 agonists. Interestingly, disease-linked Lrrc10 variants are localized to the Lrr-domain suggesting that these mutations likely alter Ca_V_1.2 interaction or binding conformation. Further studies are necessary to identify the channel interfaces responsible for binding both Lrr and the NT effector domains. One intriguing possibility is the potential binding of Lrrc10 to the carboxy-terminal domain of Ca_V_1 channels, which serves as nexus for interacting with other regulatory proteins including calmodulin and stac proteins, both of which upregulate Ca_V_1 *P*_O61,62_.

From a molecular engineering perspective, GeeC is a versatile and potentially generalizable platform to devise new Ca_V_1.2/1.3 actuators with desired properties. Our current design targets the Ca_V_β subunit to modulate channel function. As there are key differences in cytosolic domains of Ca_V_1.2 versus Ca_V_1.3, it may be possible to engineer selective nanobodies that differentially target the two closely related channels. Furthermore, at present GeeC is a constitutive enhancer of Ca_V_1 channels. However, it may be feasible to engineer GeeC such that Lrrc10 NT is recruited to the targeting domain (i.e. nanobody) through chemically-induced^63^ or optogenetic dimerizers^64^. More broadly, nanobodies have emerged as a powerful platform to probe and manipulate ion channel function. Conventionally, these nanobodies are functionalized through attachment of fluorescent reporters^65^ or enzymes that alter channel post-translational modifications to tune channel localization^48,66^. Our findings suggests that low affinity functional domains from channel interacting proteins could also be used as payloads for tuning specific aspects of channel function. This strategy may be advantages as recent studies have identified a number of short peptides that tune specific properties of Ca_V_, Na_V_ and K channels.

In all, GeeC enables targeted upregulation of Ca_V_1.2/1.3 channels in diverse physiological settings, thereby opening new avenues to develop future therapeutics and providing a convenient strategy to systematically dissect the function of these channels in a cell-type dependent manner.

## Acknowledgements

We thank members of the Calcium Signals Lab – Columbia for valuable insight and feedback on this work. This study is supported by funding from NHLBI, NINDS, and the American Heart Association. Research supported in this publication was performed in the Columbia CCTI Flow cytometry core, supported in part by the Office of the Director, National Institutes of Health under the Award S10RR027050. Images were collected and analyzed in the Confocal and Specialized Microscopy Shared Resource of the Herbert Irving Comprehensive Cancer Center at Columbia University, supported by NIH grant P30 CA013696 (National Cancer Institute). The content is solely the responsibility of the authors and does not necessarily represent the official views of the National Institutes of Health.

## Materials and Methods

### Ethical statement

All animal experimental procedures were carried out in accordance with regulations and established guidelines and were reviewed and approved by the Institutional Animal Care and Use Committee (IACUC) at Columbia University.

### Molecular Biology

We synthesized GeeC as a DNA fragment by isolating the first 53 amino acids of human LRRC10 and linking them to NbF3-P2A peptide through a GSGRSGSG sequence, flanked by NheI/EcoRI sites (Twist Bioscience). The gene fragment was ligated using T4 DNA ligase (Thermo Fisher) into a PiggyBac CMV mammalian expression vector that encoded a CFP sequence downstream of a P2A peptide. Ligates were then transformed into either XL10-Gold Ultracompetent Cells (Agilent) or DH5a Competent Cells (Thermo Fisher), plated and cultured in selective LB broth or Circlegrow Medium (MP Biomedicals). DNA was extracted and purified from cultures using either QIAprep Spin Miniprep Kit (Qiagen) or GeneJET PCR Purification Kit (Thermo Fisher). Sanger (Etion Bioscience Inc.) and Nanopore sequencing (SNPsaurus LLC) were used to verify plasmids. FRET plasmids were generated by PCR amplification of full-length human LRRC10, LRRC10 NT, and LRRC10 LRR1-CT using primers flanked by restriction sites NheI or BglII/SalI. Sequences were then ligated into a pcDNA3 plasmid coding for a downstream GSG-Cerulean (Cer) sequence immediately after restriction site.

### Adenovirus and AAV generation

Plasmids, adenovirus (pAV[Exp]-mCherry-CMV>[GEECC2.0] and pAV[Exp]-CMV>EGFP) and AAV9 (pAAV[Exp]-CMV>[GEECC2.0]:BGHpA-hPGK>mCherry:WPRE and pAAV[Exp]-CMV>EGFP:WPRE) were constructed and packaged by VectorBuilder. GeeC adenovirus vector ID is VB220916-1205wzd. GeeC AAV vector ID is VB230125-1077nkn. GFP adenovirus vector ID is VB010000-9299hac. GFP AAV vector ID is VB010000-9394npt. CFP-P2A-NbF3 adenovirus was as in previous studies.

### HEK 293 Cell Culture and Transfection

HEK293 cells (American Type Culture Collection, catalog no. CRL1573) were cultured on glass coverslips in 60-mm dishes and transfected using a calcium phosphate method. For whole-cell electrophysiology, we applied 4 µg of cDNA encoding rabbit a_1C_ subunit (NM_001136522) along with 4 µg of human β_2b_ subunit (AF285239.2) and 4 µg of rat brain a_2_d (NM_012919.2). We co-transfected 4 µg of either GeeC or NbF3 in experiments that required so. To enhance expression, cDNA for simian virus 40 T antigen (0.5 mg) was co-transfected. Electrophysiology recordings were done at room temperature 24 hours after transfection. For single-channel electrophysiology, we applied 1 µg of a_1c_, b_2b_ and a_2_d, along 1 µg of either GeeC or NbF3.

### Cardiomyocyte Isolation, Culture and Infection

Mice ventricular myocytes were isolated from 8- to 12-week-old nontransgenic (C57BL/6) mice by enzymatic digestion using a Langendorff perfusion apparatus. Both male and female mice were used to obtain isolated cardiomyocytes, and experimental outcomes were unaffected by sex. Cardiomyocytes were resuspended in perfusion solution with 5% fetal bovine serum (General Electric). CaCl_2_ was added over 30 minutes to a final concentration of 1 mM. Cells were plated on glass coverslips precoated with laminin (Corning). Plating medium contained minimal essential medium (MEM) with Earle’s salts and L-glutamine, 5% fetal bovine serum, 1% penicillin–streptomycin and 10 mM 2,3-butanedione monoxime (BDM). Cardiomyocytes were allowed to precipitate and adhere for 2 h before changing to ‘maintenance’ medium, which contained MEM with Earle’s salts, L-glutamine, 1% penicillin–streptomycin, bovine serum albumin (0.5 mg ml^−1^), 10 mM BDM, 1% insulin– transferrin–selenium, 5 mM creatine, 5 mM taurine, 2 mM L-carnitine and 25 μM blebbistatin. For deployment of GeeC, NbF3, or GFP, 1–2 μl of adenovirus were resuspended in the culture medium at least 2 hours after cell plating. Maintenance medium was replaced after 12-20 h to reduce potential adenoviral toxicity. Viral transduction efficacy was assessed 24–48 h post-infection. Only non-contracting, rod-shaped cardiomyocytes with clear striations were used for electrophysiology.

### Isolated Rat Hippocampal Neuron Culture and Infection

Culturing kits for rat hippocampi at embryonic age 18 were procured from Transetyx Tissue by BrainBits (Tranetyx Inc). Neurons were isolated from hippocampi using papain in Ca^2+^-free medium (2 mg/mL) and gentle mechanical dissociation. Cells were plated in 12-well plates containing glass cover slips precoated with poly-D-lysine (50 μg/mL) at a final concentration of 16,000 cells/cm^2^. Neurons were cultured initially in NbActiv1 culture medium (BrainBits), and switched to Neurobasal/B27/Glutamax ‘maintenance’ medium (Gibco) 96 h after plating. Medium changes were done subsequently every 72 h. GeeC and GFP were adenovirally-transduced on culture day 10 by resuspending 0.5-1 uL of adenovirus in culture medium; medium was replaced 12-20 h after infection to reduce toxicity. Transduction efficiency was evaluated 72-96 h post-infection.

### Whole-cell Electrophysiology

Whole-cell voltage-clamp recordings for HEK293 were collected at room temperature using an Axopatch 200B amplifier (Axon Instruments). Glass pipettes were pulled with a horizontal puller (P97; Sutter Instruments Co.) and fire-polished (Microforge, Narishige, Tokyo, Japan), resulting in 2–5-megaOhm resistances, before series resistance compensation of 70%. Internal solutions contained 135 mM CsMeSO_3_, 5mM CsCl_2_, 1mM MgCl_2_, 4mM MgATP, 10 mM HEPES, 10 mM BAPTA, adjusted to 290-295 mOsm with CsMeSO_3_ and pH 7.4 with CsOH. The external solutions contained 140 mM TEA-MeSO_3_, 10 mM HEPES (pH 7.4), and 5 mM CaCl_2_, 5 mM BaCl_2_ or 40 mM BaCl_2_. Solutions were adjusted to 300 mOsm with TEA-MeSO_3_ and pH 7.4 with TEA-OH. In HEK293 cells, peak currents were determined by measuring steady state currents after 15 ms test pulses from -50 to +90 mV. In cardiomyocytes, currents were elicited by 300 ms depolarization at -50 to +50 mV. Custom MATLAB (Mathworks) software was used to determine peak currents.

### Flow-cytometric FRET 2-hybrid assay

HEK293T cells were cultured in 12-well plates and transfected with polyethylenimine or FuGENE 4K (Promega). LRRC10-Cer and Holo-CaV1.2 venus (Ven)-tagged cDNA pairs (20 ng and 2 μg, respectively) were mixed in serum-free DMEM media. FRET experiments were performed 3-5 days post-transfection. Cycloheximide was added to wells at a final concentration of 100 μM 2 hours before flow-cytometric evaluation to halt new fluorophore synthesis and allow for maturation of existing fluorophores. FRET measurements were done using LSR II (BD Biosciences) flow cytometer, equipped with 405-nm, 488-nm and 633-nm lasers for excitation and 18 different emission channels. Analysis was performed as described^36^.

### Single-channel electrophysiology

Pipettes were pulled from ultra-thick-walled borosilicate glass (BF200-116-10, Sutter Instruments) as described in previous section, resulting in resistances of 5-10 megaOhm. To reduce noise, pipettes were distally coated with Sylgard (Dow Corning). External solution contained 132 K^+^-glutamate, 5 KCl, 5 NaCl, 3 MgCl_2_, 2 EGTA, 10 glucose, 20 HEPES, at 300 mOsm adjusted with glucose, and pH 7.4 adjusted with NaOH. Internal solution contained 140 mM TEA-MeSO_3_, 10 mM HEPES (pH 7.4), and 40 mM BaCl_2_, adjusted to 300 mosM with TEA-MeSO_3_ and pH 7.4 with TEA-OH. Solutions were chosen based on previous studies^61^. Single channel opening were elicited during a 200 ms voltage ramp from -80 to +60 mV. For each patch, 100-200 sweeps were recorded. Number of channels per recording were then determined by obtaining additional 80-150 sweeps after addition of Bay K 8644 to the external solution (final concentration 9 uM). Patches containing up to 5 channels were analyzed.

### Total and Surface Calcium Channel Flow-Cytometric Assay

We evaluated total and surface Ca^2+^ channel populations using flow cytometry in live, HEK293 cells as in previous studies^51^. Briefly, transfected HEK293 cells cultured in 12-well plates were gently washed with ice cold PBS containing Ca^2+^ and Mg^2+^ (in mM: 0.9 CaCl_2_, 0.49 MgCl_2_, pH 7.4) 72-96 hours post transfection. Cells were then blocked for 30 with DMEM containing 3% bovine serum albumin at 4°C, followed by 1 hour incubation in DMEM/3% bovine serum albumin containing 1 μM AlexaFluor 647 conjugated α-bungarotoxin (Life Technologies) at 4°C. HEK293 cells were then rinsed thrice with PBS containing Ca^2+^ and Mg^2+^. Cells were harvested in Ca^2+^-free PBS, and evaluated by flow cytometry using a BD Fortessa Cell Analyzer (BD Biosciences). Data was analyzed using FlowJo flow cytometry analysis software (BD Biosciences).

### Isolated Neuron Stimulation

All experiments were performed on culture day 14-16. Following a previously validated protocol, neurons were pretreated with 1 μM TTX, 10 μM APV, and 10 μM NBQX resuspended into culture medium to prevent depolarization and block glutamate-driven neurotransmission. After 4 hours, glass coverslips were transferred to a resting solution containing 150 mM NaCl, 4 mM KCl, 2 mM MgCl_2_, 2 mM CaCl_2_, 10 mM HEPES, 10 mM glucose at pH 7.4 (balanced with NaOH). Neurons were stimulated for 3 minutes by transferring coverslips into a solution containing 114 mM NaCl, 40 mM KCL, 2 mM MgCl_2_, 2 mM CaCl_2_, 10 mM HEPES, 10 mM glucose at pH 7.4 (balanced with KOH). Resting and stimulation solutions also contained 1 μM TTX, 10 μM APV and 10 μM NBQX^52^.

### Immunocytochemistry

We fixed isolated neurons using ice-cold 4% paraformaldehyde in PBS with Ca^2+^ and Mg^2+^, 4% sucrose and 20 mM EGTA. Cells were then permeabilized with 0.1% Triton X-100 (Sigma Aldrich) and blocked for 1 hour with 10% HyClone fetal bovine serum (General Electric) in PBS. Neurons were then incubated overnight at 4°C in 1% bovine serum albumin in PBS with pCREB (Ser133) (87G3) Rabbit mAb (1:333, Cell Signaling, Cat No. #9198). Neurons were then thrice rinsed with PBS, followed by 1 hour incubation with secondary antibody (1mg/mL anti-Rabbit Alexa Fluor 647, ThermoFisher) and thrice rinsing with PBS again. Coverslips were then mounted using DAPI-containing mounting solution and stored at 4°C^67,68^. *Stereotaxic Surgery.* Twelve-to fourteen-week-old C57BL/6 male mice (Jackson Laboratories, Bar Harbor, Maine) were placed on a stereotaxic surgical apparatus unit (David Kopf Instruments, Tujunga, CA). A midline incision was made on the scalp, and the head was leveled based on bregma and lambda. Mice were injected unilaterally in the dorsal hippocampus (n = 2) and the mPFC (n = 3) with two different viruses using a 2.5µl 30-gauge Hamilton syringe. The right hemisphere received 400nl injection of AAV9 – pAAV[Exp]-CMV[GEECC 2.0]-mCherry and the left hemisphere received 400nl injections of AAV9 – pAAV[Exp]-CMV-EGFP control virus for both brain regions at 0.1µl/min. Coordinates for the PFC (AP= 2.0, DV= -2.3, ML= ±0.3) and dorsal HPC (AP= -1.85, DV= -1.55, ML= ±1.10) were based on the Allen Brain Atlas. The needle was left in place for an additional 5 minutes after the infusion to ensure that the virus was completely delivered before switching to the needle with the second virus. After 2.5 weeks of recovery, mice were given a ketamine/xylazine cocktail at 10mg/kg and perfused with 1x PBS (pH 7.4) with heparin, followed by an infusion of 4% paraformaldehyde (pH 7.4). Brains were postfixed overnight in 4% PFA at 4°C and were sectioned the next day using a Leica vibratome at 40-micron thickness.

### Immunohistochemistry

Brain sections were evaluated for expression of mCherry for confirmation of AAV9-GeeC and GFP for AAV9-GFP viral infection. After confirmation of expression, tissue sections were washed 3x for 10 minutes in 1X PBS. Tissue was then blocked in a solution of 1x PBS/3% NGS/0.1% TritonX for 1 hour at room temperature, followed by 3X washes for 10 minutes in 1X PBS. Phospho-CREB (pCREB) (Ser133) (87G3) Rabbit mAb (Cell Signaling, Cat No. #9198) was diluted in 1X PBS/1% BSA/0.1% TritonX (1:1000) followed by overnight incubation at 4°C. The following day, tissue was rinsed 3X in 1X PBS for 10 minutes, and then incubated in Donkey anti-Rabbit IgG (H+L) Highly Cross-Adsorbed Secondary Antibody, Alexa Fluor™ 647 (1:500, Thermo Fisher, Cat No. A-31573). Tissue was rinsed 3X in 1X PBS for 10 minutes and then mounted on glass slides and coverslipped with ProLong™ Glass Antifade Mountant with NucBlue™ Stain (Invitrogen, Cat No. P36981). Slides were left to rest in the dark overnight before imaging.

### Image acquisition and analysis

We imaged fixed cultured rat neurons using a 60X (1.3 NA) oil objective on a Nikon Ti Eclipse inverted microscope equipped with a Yokogawa CSU-X1 confocal spinning disk and an Andor Zyla sCMOS camera. ImageJ (NIH) was used to quantify signal intensity, and we selected for analysis only neurons expressing either mCherry or GFP; uninfected controls were selected using DAPI signal. In each field of view, we selected a cell-free region of interest as signal background, and we subtracted its mean intensity from all regions of interest for each color channel. Analysis of nuclear pCREB was done by manually drawing the nuclear regions while viewing only DAPI, GFP and RFP channels, but blinded to the pCREB color channel (647 nm). We then quantified background-subtracted mean intensity and normalized it to unstimulated conditions^68^. Brain slice images were acquired for the mCherry, GFP, and Cy5 (detection of phospho-CREB) channels using a Zeiss LSM 880 Laser Scanning Confocal Microscope. All channels, laser intensities, gains and detection ranges were kept consistent between scans. GFP [488 laser(Ex. 488, Em. 525) with detection range 500-550] was used to visualize the cells infected with the AAV9-GFP reporter control, DSRed [561 laser (Ex. 561, Em. 603, detection range 580-625)] was used to image cells infected with AAV9-GeeC-mCherry, and the Cy5 channel [633 laser (Ex. 633, Em. 695, detection range 640-750)] was used to see expression of phospho-CREB after immunohistochemistry. Images were taken as a z-Stack, with bidirectional X and 4 frame averaging. The MBS 488/561/633 filter was used. After acquisition at 40x and 20x magnification, images were processed for quantification using Fiji/ImageJ. One z-stack from each scan was chosen and placed through automatic thresholding of the GeeC/GFP channels. Using Analyze Particles, automatic ROIs were made around the soma of each GeeC/GFP tagged neuron and added to the ROI manager. Alongside, automatic ROIs were created for the background to use as a normalizer for the corrected total cell fluorescence (CTCF). Area, integrated density and mean gray values were calculated for all preset ROIs in the respective pCREB/Cy5 channel stack. CTCF was calculated using Integrated Density – (Area * Mean background grey value), to normalize pCREB intensity values for the respective background around each ROI. An unpaired t-test was performed using GraphPad PRISM to compare CTCF of pCREB in GeeC and GFP tagged neurons. Results were considered significant with a p-value < 0.05.

## References

1. Hille, B. (2001). Ionic channels of excitable membranes, 3rd Edition (Sinauer Associates).

2. Zamponi, G.W., Striessnig, J., Koschak, A., and Dolphin, A.C. (2015). The Physiology, Pathology, and Pharmacology of Voltage-Gated Calcium Channels and Their Future Therapeutic Potential. Pharmacological reviews 67, 821–870. 10.1124/pr.114.009654.

3. Hofmann, F., Flockerzi, V., Kahl, S., and Wegener, J.W. (2014). L-type CaV1.2 calcium channels: from in vitro findings to in vivo function. Physiol Rev 94, 303–326. 10.1152/physrev.00016.2013.

4. Bers, D.M. (2002). Cardiac excitation-contraction coupling. Nature 415, 198–205. 10.1038/415198a.

5. Papa, A., Zakharov, S.I., Katchman, A.N., Kushner, J.S., Chen, B.X., Yang, L., Liu, G., Jimenez, A.S., Eisert, R.J., Bradshaw, G.A., et al. (2022). Rad regulation of Ca(V)1.2 channels controls cardiac fight-or-flight response. Nat Cardiovasc Res 1, 1022–1038. 10.1038/s44161-022-00157-y.

6. Vivas, O., Moreno, C.M., Santana, L.F., and Hille, B. (2017). Proximal clustering between BK and Ca(V)1.3 channels promotes functional coupling and BK channel activation at low voltage. eLife 6. 10.7554/eLife.28029.

7. Marrion, N.V., and Tavalin, S.J. (1998). Selective activation of Ca2+-activated K+ channels by co-localized Ca2+ channels in hippocampal neurons. Nature 395, 900–905. 10.1038/27674.

8. Dolmetsch, R. (2003). Excitation-transcription coupling: signaling by ion channels to the nucleus. Sci STKE 2003, PE4. 10.1126/stke.2003.166.pe4.

9. Wheeler, D.G., Groth, R.D., Ma, H., Barrett, C.F., Owen, S.F., Safa, P., and Tsien, R.W. (2012). Ca(V)1 and Ca(V)2 channels engage distinct modes of Ca(2+) signaling to control CREB-dependent gene expression. Cell 149, 1112–1124. 10.1016/j.cell.2012.03.041.

10. Dolmetsch, R.E., Pajvani, U., Fife, K., Spotts, J.M., and Greenberg, M.E. (2001). Signaling to the nucleus by an L-type calcium channel-calmodulin complex through the MAP kinase pathway. Science 294, 333–339.

11. West, A.E., Chen, W.G., Dalva, M.B., Dolmetsch, R.E., Kornhauser, J.M., Shaywitz, A.J., Takasu, M.A., Tao, X., and Greenberg, M.E. (2001). Calcium regulation of neuronal gene expression. Proc Natl Acad Sci U S A 98, 11024–11031.

12. Schulla, V., Renstrom, E., Feil, R., Feil, S., Franklin, I., Gjinovci, A., Jing, X.J., Laux, D., Lundquist, I., Magnuson, M.A., et al. (2003). Impaired insulin secretion and glucose tolerance in beta cell-selective Ca(v)1.2 Ca2+ channel null mice. The EMBO journal 22, 3844–3854. 10.1093/emboj/cdg389.

13. Marcantoni, A., Vandael, D.H., Mahapatra, S., Carabelli, V., Sinnegger-Brauns, M.J., Striessnig, J., and Carbone, E. (2010). Loss of Cav1.3 channels reveals the critical role of L-type and BK channel coupling in pacemaking mouse adrenal chromaffin cells. J Neurosci 30, 491–504. 10.1523/JNEUROSCI.4961-09.2010.

14. Hill-Eubanks, D.C., Werner, M.E., Heppner, T.J., and Nelson, M.T. (2011). Calcium signaling in smooth muscle. Cold Spring Harbor perspectives in biology 3, a004549. 10.1101/cshperspect.a004549.

15. Brandt, A., Striessnig, J., and Moser, T. (2003). CaV1.3 channels are essential for development and presynaptic activity of cochlear inner hair cells. J Neurosci 23, 10832–10840. 10.1523/JNEUROSCI.23-34-10832.2003.

16. Pitt, G.S., Matsui, M., and Cao, C. (2021). Voltage-Gated Calcium Channels in Nonexcitable Tissues. Annu Rev Physiol 83, 183–203. 10.1146/annurev-physiol-031620-091043.

17. Ortner, N.J., Kaserer, T., Copeland, J.N., and Striessnig, J. (2020). De novo CACNA1D Ca(2+) channelopathies: clinical phenotypes and molecular mechanism. Pflugers Arch 472, 755–773. 10.1007/s00424-020-02418-w.

18. Herold, K.G., Hussey, J.W., and Dick, I.E. (2023). CACNA1C-Related Channelopathies. Handb Exp Pharmacol 279, 159–181. 10.1007/164_2022_624.

19. Liao, P., and Soong, T.W. (2010). CaV1.2 channelopathies: from arrhythmias to autism, bipolar disorder, and immunodeficiency. Pflugers Arch 460, 353–359. 10.1007/s00424-009-0753-0.

20. Splawski, I., Timothy, K.W., Sharpe, L.M., Decher, N., Kumar, P., Bloise, R., Napolitano, C., Schwartz, P.J., Joseph, R.M., Condouris, K., et al. (2004). Ca(V)1.2 calcium channel dysfunction causes a multisystem disorder including arrhythmia and autism. Cell 119, 19–31.

21. Endres, D., Decher, N., Rohr, I., Vowinkel, K., Domschke, K., Komlosi, K., Tzschach, A., Glaser, B., Schiele, M.A., Runge, K., et al. (2020). New Cav1.2 Channelopathy with High-Functioning Autism, Affective Disorder, Severe Dental Enamel Defects, a Short QT Interval, and a Novel CACNA1C Loss-Of-Function Mutation. International journal of molecular sciences 21. 10.3390/ijms21228611.

22. Rodan, L.H., Spillmann, R.C., Kurata, H.T., Lamothe, S.M., Maghera, J., Jamra, R.A., Alkelai, A., Antonarakis, S.E., Atallah, I., Bar-Yosef, O., et al. (2021). Phenotypic expansion of CACNA1C-associated disorders to include isolated neurological manifestations. Genet Med 23, 1922–1932. 10.1038/s41436-021-01232-8.

23. Yu, W. (2022). Reviving Cav1.2 as an attractive drug target to treat bladder dysfunction. FASEB J 36, e22118. 10.1096/fj.202101475R.

24. Hockerman, G.H., Peterson, B.Z., Johnson, B.D., and Catterall, W.A. (1997). Molecular determinants of drug binding and action on L-type calcium channels. Annual review of pharmacology and toxicology 37, 361–396. 10.1146/annurev.pharmtox.37.1.361.

25. Bourson, A., Moser, P.C., Gower, A.J., and Mir, A.K. (1989). Central and peripheral effects of the dihydropyridine calcium channel activator BAY K 8644 in the rat. European journal of pharmacology 160, 339–347. 10.1016/0014-2999(89)90089-7.

26. Shelton, R.C., Grebb, J.A., and Freed, W.J. (1987). Induction of seizures in mice by intracerebroventricular administration of the calcium channel agonist BAY k 8644. Brain research 402, 399–402. 10.1016/0006-8993(87)90054-0.

27. Jinnah, H.A., Yitta, S., Drew, T., Kim, B.S., Visser, J.E., and Rothstein, J.D. (1999). Calcium channel activation and self-biting in mice. Proc Natl Acad Sci U S A 96, 15228–15232. 10.1073/pnas.96.26.15228.

28. Chen, H., Vandorpe, D.H., Xie, X., Alper, S.L., Zeidel, M.L., and Yu, W. (2020). Disruption of Cav1.2-mediated signaling is a pathway for ketamine-induced pathology. Nature communications 11, 4328. 10.1038/s41467-020-18167-4.

29. Kabir, Z.D., Martinez-Rivera, A., and Rajadhyaksha, A.M. (2017). From Gene to Behavior: L-Type Calcium Channel Mechanisms Underlying Neuropsychiatric Symptoms. Neurotherapeutics 14, 588–613. 10.1007/s13311-017-0532-0.

30. Heyes, S., Pratt, W.S., Rees, E., Dahimene, S., Ferron, L., Owen, M.J., and Dolphin, A.C. (2015). Genetic disruption of voltage-gated calcium channels in psychiatric and neurological disorders. Prog Neurobiol 134, 36–54. 10.1016/j.pneurobio.2015.09.002.

31. International Schizophrenia, C., Purcell, S.M., Wray, N.R., Stone, J.L., Visscher, P.M., O’Donovan, M.C., Sullivan, P.F., and Sklar, P. (2009). Common polygenic variation contributes to risk of schizophrenia and bipolar disorder. Nature 460, 748–752. 10.1038/nature08185.

32. Cross-Disorder Group of the Psychiatric Genomics, C. (2013). Identification of risk loci with shared effects on five major psychiatric disorders: a genome-wide analysis. Lancet 381, 1371-1379. 10.1016/S0140-6736(12)62129-1.

33. Ament, S.A., Szelinger, S., Glusman, G., Ashworth, J., Hou, L., Akula, N., Shekhtman, T., Badner, J.A., Brunkow, M.E., Mauldin, D.E., et al. (2015). Rare variants in neuronal excitability genes influence risk for bipolar disorder. Proc Natl Acad Sci U S A 112, 3576–3581. 10.1073/pnas.1424958112.

34. Pinggera, A., and Striessnig, J. (2016). Ca(v) 1.3 (CACNA1D) L-type Ca(2+) channel dysfunction in CNS disorders. J Physiol 594, 5839–5849. 10.1113/JP270672.

35. Catterall, W.A., Lenaeus, M.J., and Gamal El-Din, T.M. (2020). Structure and Pharmacology of Voltage-Gated Sodium and Calcium Channels. Annual review of pharmacology and toxicology 60, 133–154. 10.1146/annurev-pharmtox-010818-021757.

36. Liu, G., Papa, A., Katchman, A.N., Zakharov, S.I., Roybal, D., Hennessey, J.A., Kushner, J., Yang, L., Chen, B.X., Kushnir, A., et al. (2020). Mechanism of adrenergic CaV1.2 stimulation revealed by proximity proteomics. Nature 577, 695–700. 10.1038/s41586-020-1947-z.

37. Ben-Johny, M., and Yue, D.T. (2014). Calmodulin regulation (calmodulation) of voltage-gated calcium channels. J Gen Physiol 143, 679–692. 10.1085/jgp.201311153.

38. Woon, M.T., Long, P.A., Reilly, L., Evans, J.M., Keefe, A.M., Lea, M.R., Beglinger, C.J., Balijepalli, R.C., Lee, Y., Olson, T.M., and Kamp, T.J. (2018). Pediatric Dilated Cardiomyopathy-Associated LRRC10 (Leucine-Rich Repeat-Containing 10) Variant Reveals LRRC10 as an Auxiliary Subunit of Cardiac L-Type Ca(2+) Channels. Journal of the American Heart Association 7. 10.1161/JAHA.117.006428.

39. Manuylov, N.L., Manuylova, E., Avdoshina, V., and Tevosian, S. (2008). Serdin1/Lrrc10 is dispensable for mouse development. Genesis 46, 441–446. 10.1002/dvg.20422.

40. Nguyen, P.D., Gooijers, I., Campostrini, G., Verkerk, A.O., Honkoop, H., Bouwman, M., de Bakker, D.E.M., Koopmans, T., Vink, A., Lamers, G.E.M., et al. (2023). Interplay between calcium and sarcomeres directs cardiomyocyte maturation during regeneration. Science 380, 758–764. 10.1126/science.abo6718.

41. Stockdale, W.T., Lemieux, M.E., Killen, A.C., Zhao, J., Hu, Z., Riepsaame, J., Hamilton, N., Kudoh, T., Riley, P.R., van Aerle, R., et al. (2018). Heart Regeneration in the Mexican Cavefish. Cell Rep 25, 1997–2007 e1997. 10.1016/j.celrep.2018.10.072.

42. Kim, K.H., Antkiewicz, D.S., Yan, L., Eliceiri, K.W., Heideman, W., Peterson, R.E., and Lee, Y. (2007). Lrrc10 is required for early heart development and function in zebrafish. Developmental biology 308, 494–506. 10.1016/j.ydbio.2007.06.005.

43. Salamon, R.J., McKeon, M.C., Bae, J., Zhang, X., Paltzer, W.G., Wanless, K.N., Schuett, A.R., Nuttall, D.J., Nemr, S.A., Sridharan, R., et al. (2023). LRRC10 regulates mammalian cardiomyocyte cell cycle during heart regeneration. NPJ Regen Med 8, 39. 10.1038/s41536-023-00316-0.

44. Brody, M.J., Feng, L., Grimes, A.C., Hacker, T.A., Olson, T.M., Kamp, T.J., Balijepalli, R.C., and Lee, Y. (2016). LRRC10 is required to maintain cardiac function in response to pressure overload. American journal of physiology. Heart and circulatory physiology 310, H269–278. 10.1152/ajpheart.00717.2014.

45. Matsushima, N., Miyashita, H., Mikami, T., and Kuroki, Y. (2010). A nested leucine rich repeat (LRR) domain: the precursor of LRRs is a ten or eleven residue motif. BMC Microbiol 10, 235. 10.1186/1471-2180-10-235.

46. Enkhbayar, P., Kamiya, M., Osaki, M., Matsumoto, T., and Matsushima, N. (2004). Structural principles of leucine-rich repeat (LRR) proteins. Proteins 54, 394–403. 10.1002/prot.10605.

47. Matsushima, N., Takatsuka, S., Miyashita, H., and Kretsinger, R.H. (2019). Leucine Rich Repeat Proteins: Sequences, Mutations, Structures and Diseases. Protein Pept Lett 26, 108–131. 10.2174/0929866526666181208170027.

48. Morgenstern, T.J., Park, J., Fan, Q.R., and Colecraft, H.M. (2019). A potent voltage-gated calcium channel inhibitor engineered from a nanobody targeted to auxiliary Ca(V)beta subunits. Elife 8. 10.7554/eLife.49253.

49. Buraei, Z., and Yang, J. (2010). The beta subunit of voltage-gated Ca2+ channels. Physiol Rev 90, 1461–1506. 10.1152/physrev.00057.2009.

50. Perez-Reyes, E. (2003). Molecular physiology of low-voltage-activated t-type calcium channels. Physiol Rev 83, 117–161. 10.1152/physrev.00018.2002.

51. Yang, T., Xu, X., Kernan, T., Wu, V., and Colecraft, H.M. (2010). Rem, a member of the RGK GTPases, inhibits recombinant CaV1.2 channels using multiple mechanisms that require distinct conformations of the GTPase. J Physiol 588, 1665–1681.

52. Li, B., Tadross, M.R., and Tsien, R.W. (2016). Sequential ionic and conformational signaling by calcium channels drives neuronal gene expression. Science 351, 863–867. 10.1126/science.aad3647.

53. Khan, I.F., Hirata, R.K., and Russell, D.W. (2011). AAV-mediated gene targeting methods for human cells. Nature protocols 6, 482–501. 10.1038/nprot.2011.301.

54. Kamijo, S., Ishii, Y., Horigane, S.I., Suzuki, K., Ohkura, M., Nakai, J., Fujii, H., Takemoto-Kimura, S., and Bito, H. (2018). A Critical Neurodevelopmental Role for L-Type Voltage-Gated Calcium Channels in Neurite Extension and Radial Migration. J Neurosci 38, 5551–5566. 10.1523/JNEUROSCI.2357-17.2018.

55. Muth, J.N., Bodi, I., Lewis, W., Varadi, G., and Schwartz, A. (2001). A Ca(2+)-dependent transgenic model of cardiac hypertrophy: A role for protein kinase Calpha. Circulation 103, 140–147. 10.1161/01.cir.103.1.140.

56. Ahern, B.M., Levitan, B.M., Veeranki, S., Shah, M., Ali, N., Sebastian, A., Su, W., Gong, M.C., Li, J., Stelzer, J.E., et al. (2019). Myocardial-restricted ablation of the GTPase RAD results in a pro-adaptive heart response in mice. J Biol Chem 294, 10913–10927. 10.1074/jbc.RA119.008782.

57. Brody, M.J., and Lee, Y. (2016). The Role of Leucine-Rich Repeat Containing Protein 10 (LRRC10) in Dilated Cardiomyopathy. Front Physiol 7, 337. 10.3389/fphys.2016.00337.

58. Brody, M.J., Hacker, T.A., Patel, J.R., Feng, L., Sadoshima, J., Tevosian, S.G., Balijepalli, R.C., Moss, R.L., and Lee, Y. (2012). Ablation of the cardiac-specific gene leucine-rich repeat containing 10 (Lrrc10) results in dilated cardiomyopathy. PloS one 7, e51621. 10.1371/journal.pone.0051621.

59. Qu, X.K., Yuan, F., Li, R.G., Xu, L., Jing, W.F., Liu, H., Xu, Y.J., Zhang, M., Liu, X., Fang, W.Y., et al. (2015). Prevalence and spectrum of LRRC10 mutations associated with idiopathic dilated cardiomyopathy. Mol Med Rep 12, 3718–3724. 10.3892/mmr.2015.3843.

60. Huang, L., Tang, S., Chen, Y., Zhang, L., Yin, K., Wu, Y., Zheng, J., Wu, Q., Makielski, J.C., and Cheng, J. (2017). Molecular pathological study on LRRC10 in sudden unexplained nocturnal death syndrome in the Chinese Han population. International journal of legal medicine 131, 621–628. 10.1007/s00414-016-1516-z.

61. Adams, P.J., Ben-Johny, M., Dick, I.E., Inoue, T., and Yue, D.T. (2014). Apocalmodulin itself promotes ion channel opening and Ca(2+) regulation. Cell 159, 608–622. 10.1016/j.cell.2014.09.047.

62. Niu, J., Dick, I.E., Yang, W., Bamgboye, M.A., Yue, D.T., Tomaselli, G., Inoue, T., and Ben-Johny, M. (2018). Allosteric regulators selectively prevent Ca(2+)-feedback of Ca(V) and Na(V) channels. eLife 7. 10.7554/eLife.35222.

63. Stanton, B.Z., Chory, E.J., and Crabtree, G.R. (2018). Chemically induced proximity in biology and medicine. Science 359. 10.1126/science.aao5902.

64. Niu, J., Ben Johny, M., Dick, I.E., and Inoue, T. (2016). Following Optogenetic Dimerizers and Quantitative Prospects. Biophys J 111, 1132–1140. 10.1016/j.bpj.2016.07.040.

65. Gross, G.G., Junge, J.A., Mora, R.J., Kwon, H.B., Olson, C.A., Takahashi, T.T., Liman, E.R., Ellis-Davies, G.C., McGee, A.W., Sabatini, B.L., et al. (2013). Recombinant probes for visualizing endogenous synaptic proteins in living neurons. Neuron 78, 971–985. 10.1016/j.neuron.2013.04.017.

66. Kanner, S., Morgenstern, T.J., and Colecraft, H.M. (2017). Sculpting ion channel functional expression with engineered ubiquitin ligases. eLife In press.

67. Deisseroth, K., Heist, E.K., and Tsien, R.W. (1998). Translocation of calmodulin to the nucleus supports CREB phosphorylation in hippocampal neurons. Nature 392, 198–202.

68. Li, B., Tadross, M.R., and Tsien, R.W. (2016). Sequential ionic and conformational signaling by calcium channels drives neuronal gene expression. Science 351, 863–867.

